# Generation of a novel constitutive smooth muscle cell-specific *Myh11*-driven Cre mouse model

**DOI:** 10.1101/2024.11.08.622724

**Authors:** Kunzhe Dong, Zhixia Bai, Xiangqin He, Lu Zhang, Guoqing Hu, Yali Yao, Chen-Leng Cai, Jiliang Zhou

## Abstract

Dysfunction in either embryonic or postnatal vascular smooth muscle cells (SMCs) significantly contributes to the progression of various cardiovascular diseases. Therefore, elucidating the molecular mechanisms governing VSMC development and homeostasis is crucial. *MYH11* is the most reliable lineage gene for SMCs and has been utilized to develop tamoxifen-inducible Cre driver lines for achieving SMC-specific gene manipulation by crossing with mice carrying the lox*P*-flanked gene, particularly in adult mice. For studies involving SMCs during embryogenesis, the commonly used constitutive Cre driver is controlled by the *Tagln* (*Sm22α*) promoter. However, this Cre driver exhibits activity in multiple non-SMC populations, including cardiomyocytes and skeletal muscle precursors, introducing confounding effects. Additionally, most existing SMC-specific Cre drivers are generated using a transgenic approach, raising concerns about random site integration and variable gene copy numbers. To address these limitations, we report a novel Cre mouse model generated by knock-in (KI) of a nuclear-localized Cre recombinase into the *Myh11* gene locus using homologous recombination. We confirmed that the Cre activity precisely recapitulates endogenous *Myh11* expression by crossing with *Rosa26* mTmG or tdTomato reporter mice. Moreover, *Myh11*-driven Cre can efficiently delete the floxed allele of the transcription factor *Tead1* specifically in SMCs. The *Tead1* SMC-specific knockout mice did not exhibit an overt phenotype, thereby circumventing the embryonic lethal phenotype mediated by *Tagln*-driven Cre, as we previously reported. These findings establish this novel Cre driver line as a robust tool for tracing the *Myh11*-positive SMC lineage and manipulating gene function specifically in SMCs during embryonic development in mice.

## INTRODUCTION

Smooth muscle cells (SMCs) are the principal cellular components of the walls in most hollow organs including blood vessels and gastrointestinal (GI) tissues. During embryogenesis, SMCs originate from multiple progenitor lineages and contribute to the development of these organs. Defects in SMC development can lead to impaired vascular architecture ^1-3^, various congenital cardiovascular diseases ^4-7^, visceral myopathy ^8^ and increased susceptibility to cardiovascular diseases ^9-13^. Mature vascular SMCs are highly differentiated and express a unique set of contractile genes. In response to pathophysiological stimuli, vascular SMCs undergo phenotypic modulation from a differentiation to a dedifferentiation state, adopting phenotypes resembling diverse cell types, including stem cells, fibroblasts, macrophages, and chondrocytes ^14,15^. This process, known as phenotypic switching of SMCs, significantly contributes to the progression of various vascular diseases such as atherosclerosis and restenosis ^16^. Notably, lineage tracing studies in mice suggest that 30-70% of the cells within atherosclerotic plaques ^17-19^ and approximately 80% of the cells within injury-induced neointima ^20-22^ are derived from local vascular SMCs. Furthermore, recent genome-wide epigenetic studies reveal that GWAS-identified risk SNPs for various vascular diseases, such as coronary artery disease, blood pressure disorders, and stroke, are enriched in vascular SMC-specific open chromatin regions ^23,24^, highlighting an autonomous mechanism of vascular SMCs in vascular pathology. However, the functional roles of genes associated with SMC-driven cardiovascular diseases in vivo remain largely unknown due to a lack of a more SMC-restrictive Cre driver.

The Cre/loxP system is a powerful and widely used strategy for elucidating gene function in specific cell populations in mice ^25^. Multiple Cre driver lines have been developed to study SMC biology, primarily by utilizing promoters of *Myh11, Tagln* (also known as SM22α), or *Acta2* ^26-28^. While *Tagln* and *Acta2* show non-SMC expression patterns, *Myh11* is widely acknowledged as the most specific SMC lineage gene ^29^, making it the preferred candidate for generating SMC-specific Cre drivers ^26,27,30^. Several inducible Cre-ER^T2^ driver lines have been developed using the *Myh11* promoter, including both constitutive and inducible Cre systems ^30-32^. For embryonic SMC investigations, the commonly used Cre driver is the one developed by Herz’s laboratory, which employs a 2.8 kb fragment of the *Tagln* promoter (Jackson Laboratories # B6.Cg-Tg[Tagln-cre]1Her/J) ^33^. However, the non-specific expression of *Tagln*, especially its transient activation in cardiomyocytes during early embryonic stages, raises concerns when using this Cre driver to study SMC-specific gene effects ^26^. Furthermore, most existing SMC-specific Cre drivers are generated using transgenic strategies, which introduces additional concerns regarding random site integration and the variable number of transgene copies ^34^.

To address these limitations, we generated a novel Cre mouse model by knocking in (KI) a nuclear-localized Cre recombinase into the *Myh11* gene locus using a homologous recombination approach. We confirmed that the Cre activity precisely recapitulates endogenous *Myh11* expression in mice by breeding with *Rosa26* mTmG (membrane-targeted tandem dimer Tomato, mTomato or mT; membrane-targeted green fluorescent protein, mGFP or mG) or tdTomato reporter mice. Additionally, *Myh11*-driven Cre efficiently deletes the floxed allele of the transcription factor *Tead1* specifically in SMCs, while the *Tead1* SMC-specific KO mice did not exhibit an overt phenotype, thus circumventing the embryonic lethal phenotype mediated by *Tagln*-driven Cre, as we previously reported ^6^. These findings establish this novel driver line as a robust tool for tracing *Myh11*-positive SMC lineages and for manipulating gene function specifically in SMCs during development in mice.

## RESULTS

### Generation of a constitutive *Myh11*^*Cre/+*^ mouse line

To generate a Cre driver that constitutively expresses Cre recombinase specifically in SMCs, we inserted a NLS (nuclear localization sequence PKKKRKV from Simian Virus 40 large T-antigen)-CreER^T2^-polyA cassette immediately downstream of the start codon (ATG) of the endogenous *Myh11* gene, the most specific gene for SMC lineage ^26,35^, using an homologous recombination strategy as we recently reported ^35,36^ (**Figure 1A**). The NLS-mediated Cre nuclear location enables constitutive Cre recombination activity despite the presence of ER^T2^ cassette. This approach also ensures the SMC-restricted production of Cre under the control of *Myh11* gene promoter in both sexes of mice because the *Myh11* gene is on autosomal chromosome 16 in mice. The Cre line was maintained as heterozygotes (*Myh11*^*Cre/+*^) by crossing with wild-type (WT) mice. The proper insertion of the targeted KI cassette was verified by PCR in both male and female mice (**Figure 1B**). The DNA integrity of the insertion was further validated by Sanger sequencing (**Supplemental Figure 1A-B**). To assess Cre production, we performed Western Blotting analysis using protein lysates from SMC-enriched tissues including aorta, colon and bladder from both sexes of *Myh11*^*Cre/+*^ mice and their *Myh11*^*+/+*^ (Cre negative) WT littermates. This analysis revealed robust Cre expression in all examined tissues with comparable expression in both sexes of *Myh11*^*Cre/+*^ mice, but not in WT mice (**Figure 1C**). Furthermore, data from these Western blotting assays revealed that *Myh11*^*Cre/+*^ mice exhibit a slight but insignificant reduction in MYH11 protein expression (**Figure 1D**), suggesting that a single WT allele can largely compensate for the loss of the other KI allele as we recently described ^35^. These results collectively indicate the successful generation of a novel constitutive *Myh11*^*Cre*^ KI driver line, with negligible loss of endogenous MYH11 expression in the aorta and in SMC-enriched visceral tissues such as the colon and bladder.

**Figure 1.**
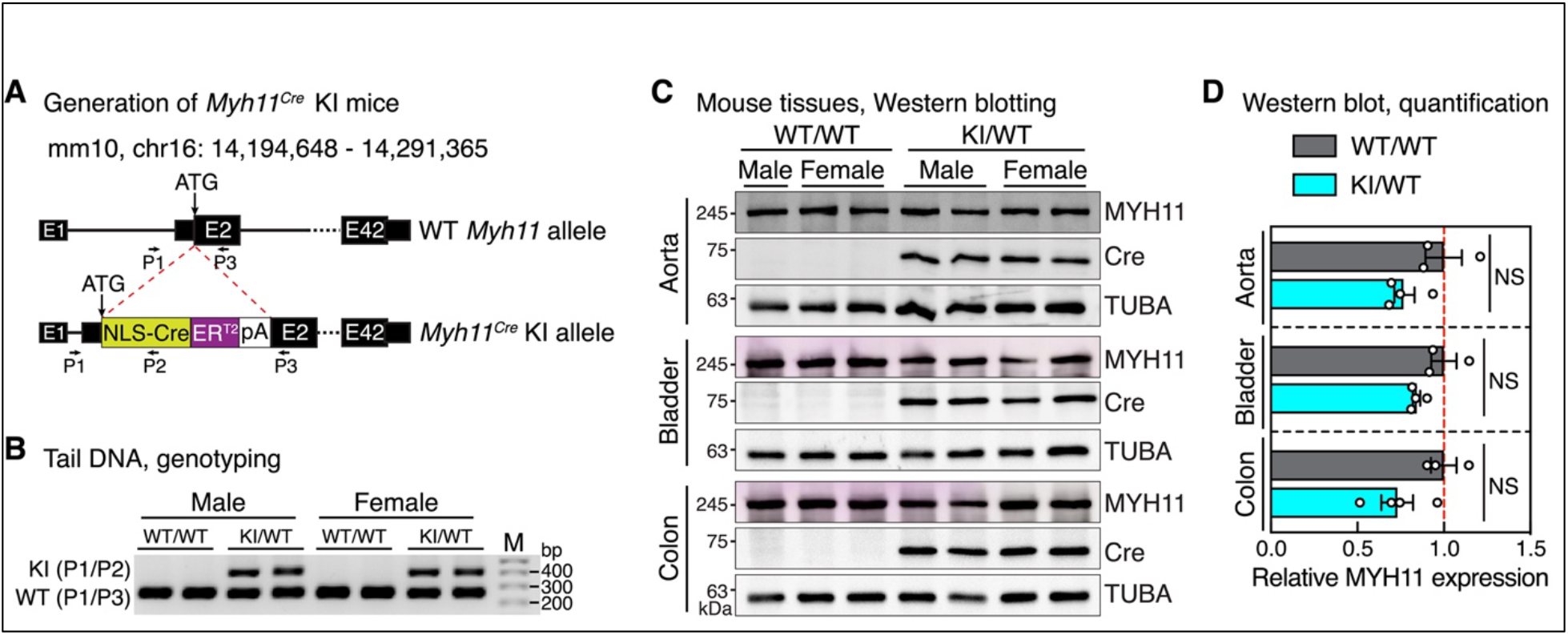
Generation of *Myh11*^*Cre*^ KI mice. **A**. Schematic showing the strategy for generating the *Myh11*^*Cre*^ mice using homologous recombination. Primers (P) P1, P2 and P3 were used for genotyping and the sequences of these primers are provided in Supplemental Figure 1. **B**. PCR analysis of tail genomic DNA showing the presence of the KI allele in both male and female mice. M: DNA marker. **C**. Western blotting analysis of CreER^T2^ and endogenous MYH11 expression in aorta, bladder and colon isolated from adult male or female WT and KI heterozygous mice. **D**. The densitometry of the Western blotting shown in panel “**C**” was quantified and the relative expression of MYH11 calculated after normalizing to the loading control TUBA protein is plotted (WT/WT was set to 1). N=3 or 4. NS, not statistically significant.

### Cre activity driven by endogenous *Myh11* gene is specific to SMCs

To assess specificity of the *Myh11*^*Cre/+*^ driver, we introduced the *Myh11*^*Cre/+*^ mice into mTmG double-fluorescence reporter mice (*mTmG*^*F/F*^) ^37^, in which most cells express tdTomato fluorescence under the ubiquitous *Rosa26* promoter (**Figure 2A-B**). In the resulting *Myh11*^*Cre/+*^*;mTmG*^*F/W*^ offspring, cells only undergoing Cre/lox*P* recombination express GFP fluorescence resulting from the removal of the mT and stop cassette. Therefore, GFP fluorescence can be used to trace *Myh11* positive SMC lineage and Cre-mediated DNA recombination events (**Figure 2A-B**). We first examined GFP expression in the cardiovascular system including heart and aorta. We found that GFP signal is specifically observed in aortic tissues and coronary arteries of *Myh11*^*Cre/+*^*;mTmG*^*F/W*^ mice, but not *Myh11*^*+/+*^*;mTmG*^*F/W*^ control mice (**Figure 2B-C**). Examination of GFP fluorescence in sections of aorta and heart further revealed that GFP signal was exclusively present in the medial layer of aortic tissues and coronary arteries where VSMCs are located, but not in tdTomato-expressing endothelial, adventitial, and cardiac muscle cells (**Figure 2D-E**).

**Figure 2.**
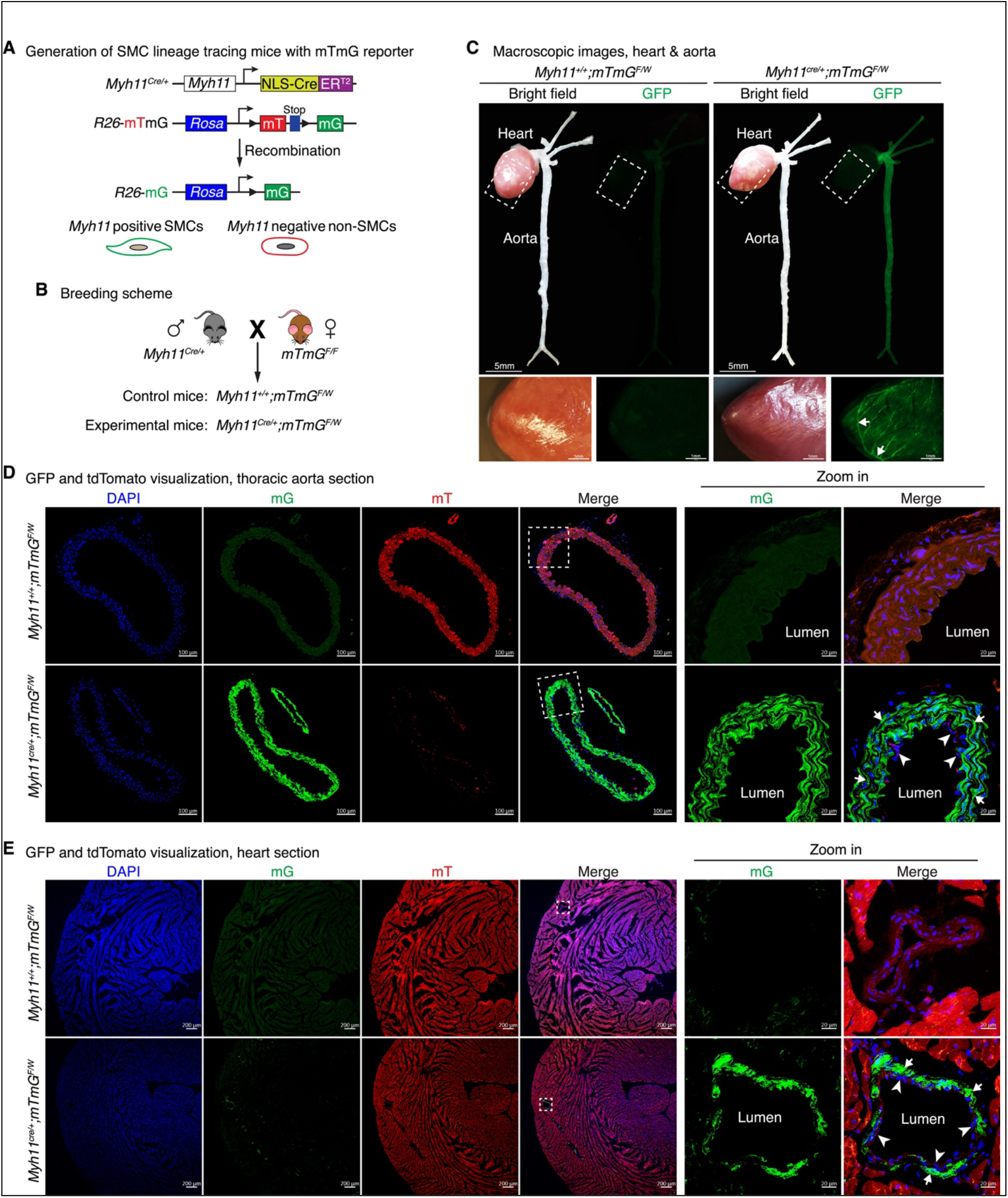
Examination of Cre activity of the *Myh11*-Cre driver in cardiovascular tissues using the mTmG dual fluorescence reporter mice. **A-B**. Strategy (**A**) and breeding scheme (**B**) for generating SMC lineage tracing mice by crossing *Myh11*^*Cre*^ mice with mTmG reporter mice. Triangles indicate loxP sites. **C**. Visualization of GFP in heart and aorta of *Myh11*^*+/+*^*;mTmG*^*F/W*^ control and *Myh11*^*Cre/+*^*;mTmG*^*F/W*^ SMC lineage tracing mice. The boxed areas in the heart are magnified on the bottom of respective panels. White arrows point to the representative coronary arteries. **D-E**. GFP and tdTomato visualization in sections of aorta **(D)** and heart **(E)** of control and SMC lineage tracing mice. Boxed areas are magnified on the right. Arrows indicate presumptive GFP positive SMCs and arrowheads point to the tdTomato positive endothelial cells in both thoracic aorta (**D**) and coronary artery (**E**) of the heart, respectively. Cell nuclei were stained with DAPI (blue).

We next extended our characterization of GFP signaling to other tissues. A gross examination of the brain, GI tract, and quadricep muscles revealed the presence of GFP exclusively in bladder, small intestine, and blood vessels of the brain and quadriceps (**Supplemental Figure 2A**). Subsequent GFP visualization of these tissue sections indicated GFP expression restricted to the presumptive SMC layers of tissues such as bladder, brain, colon, esophagus, intestine, kidney, lung, quadricep muscle, stomach, trachea, and male vas deferens (**Supplemental Figure 2B**). This observed pattern aligns completely with that of endogenous *Myh11* gene as previously reported ^29,36^. In contrast to a previous report ^38^, we did not observe GFP signal in interstitial cells of the thymus in our *Myh11-Cre*^*Cre/+*^; *mTmG*^*F/W*^ mice (**Supplemental Figure 2C**).

To further confirm whether Cre activity driven by the endogenous *Myh11* gene recapitulates *Myh11* expression, we crossed *Myh11*^*Cre/+*^ male mice with single fluorescence tdTomato reporter female mice (**Figure 3A**) and performed immunofluorescence (IF) staining for MYH11 in their offspring *Myh11*^*Cre/+*^*;tdTomato*^*F/W*^ mice. Direct visualization of the RFP signal in *Myh11*^*Cre/+*^*;tdTomato*^*F/W*^ mouse tissues revealed that RFP is specifically observed in developmental GI tissues such as the E16.5 intestine, postnatal day 0 (P0) stomach, and coronary arteries in the heart of adult mice (**Figure 3B-C**). Co-IF staining for MYH11, the endothelial cell marker PECAM1 and native tdTomato fluorescence signal demonstrated that tdTomato driven by the *Myh11* gene perfectly overlaps with the MYH11 IF signal but not with PECAM1 in the developing GI tissues and cardiovascular tissues of adult mice (**Figure 3D**).

**Figure 3.**
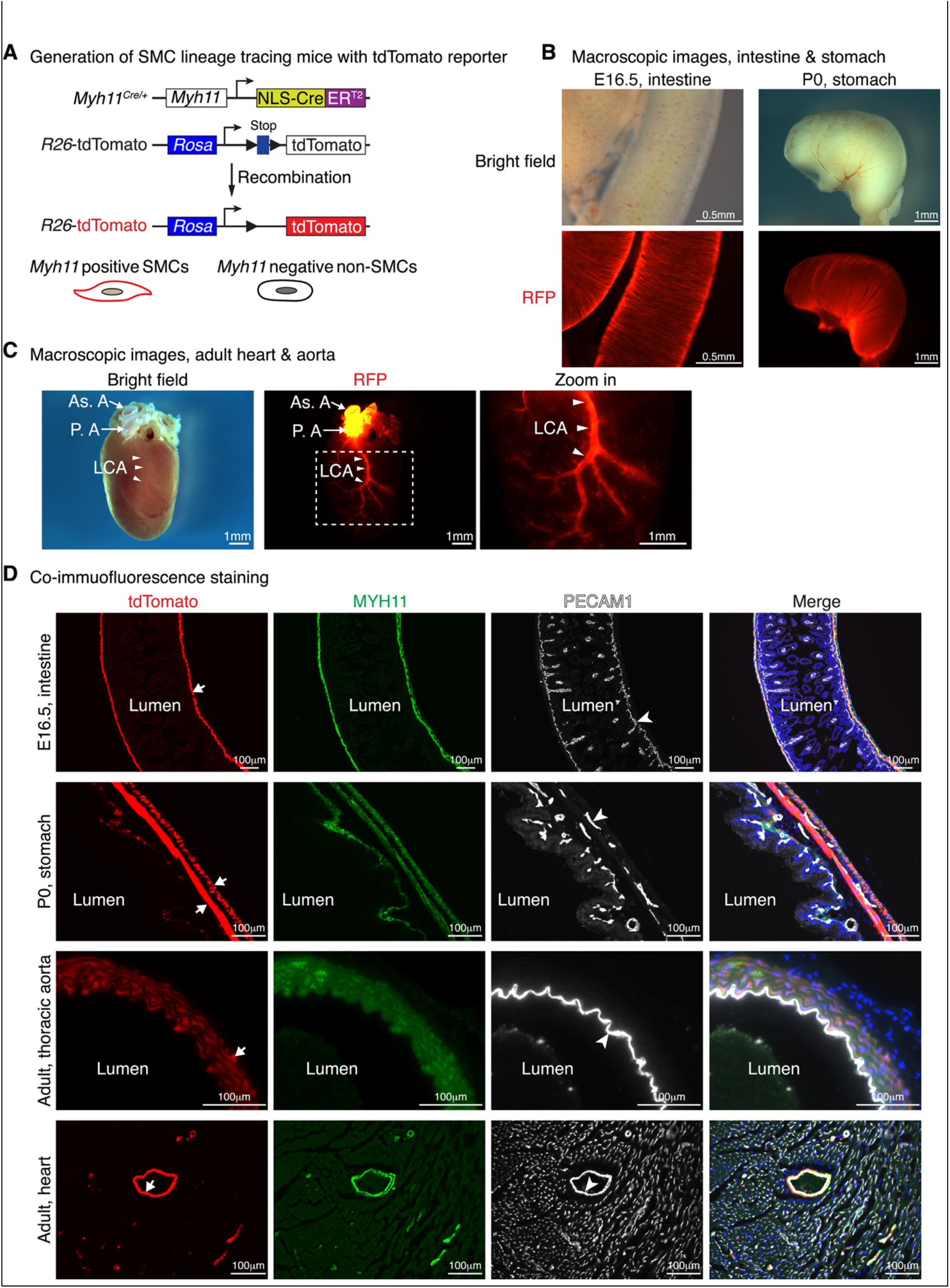
Examination of Cre activity of the *Myh11*-Cre driver using the tdTomato single fluorescence reporter mice. **A**. Schematic depiction of the strategy to generate the tdTomato single fluorescence SMC lineage tracing mice by crossing *Myh11*^*Cre/+*^ mice with tdTomato reporter mice. Triangles indicate loxP sites. **B-C**. Visualization of RFP of E16.5 intestine (left panel in “**B**”), P0 stomach (right panel in “**B**”) and the heart of *Myh11*^*Cre/+*^*;tdTomato*^*F/W*^ adult mice under bright field and RFP channels. The boxed areas in the heart (**C**) are magnified to the right showing RFP-positive left coronary artery (LCA, denoted by arrowheads). As. A: ascending aorta; P. A: pulmonary artery. **D**. Co-IF of MYH11 (green), the endothelial cell marker PECAM1 (white) and visualization of native tdTomato signal (red) in sections of tissues harvested from tdTomato SMC lineage tracing mice as indicated. Arrows indicate double positive SMCs of both tdTomato and MYH11. Arrowheads point to the PECAM1 positive endothelial cells. Cell nuclei were stained with Hoechst 33342 (blue).

Taken together, examination of these two independent lineage tracing mouse models, along with MYH11 IF, demonstrates that Cre expression driven by the endogenous *Myh11* gene is specific in SMCs, and Cre-mediated recombination selectively occurs in vascular and visceral SMCs.

### *Myh11*^*Cre/+*^ mediates the deletion of the floxed *Tead1* gene specifically in SMCs

We previously demonstrated that *Tagln*-Cre-mediated deletion of the transcription factor *Tead1* in cardiac and SMCs leads to embryonic lethality by E14.5 as a result of impaired cardiomyocyte and SMC proliferation and differentiation ^6^. To determine whether *Myh11*^*Cre*^ KI mice can efficiently delete the floxed *Tead1* allele specifically in SMCs, but not in cardiomyocytes during development, we crossed *Myh11*^*Cre*^ KI mice with floxed *Tead1* mice to generate constitutive SMC-specific *Tead1* KO mice (**Figure 4A-B**). The specific and efficient deletion of *Tead1* in aortic SMCs versus cardiomyocytes was confirmed by IF staining and Western blotting of TEAD1 in adult mice (**Figure 4C-F**). In contrast to the lethal embryonic phenotype seen in *Tagln*-Cre-mediated *Tead1* KO mice ^6^, *Myh11-Cre-mediated Tead1* KO mice are born at the expected Mendelian ratios, fertile, and live up to the full life span without an overt phenotype (data not shown), indicating *Myh11*-Cre can circumvent the lethal cardiac phenotype mediated by *Tagln*-Cre. Taken together, these data demonstrate the specificity and utility of this novel constitutive SMC-specific Cre driver for SMC lineage tracing and for examining the functional role of genes of interest during and following mouse embryogenesis.

**Figure 4.**
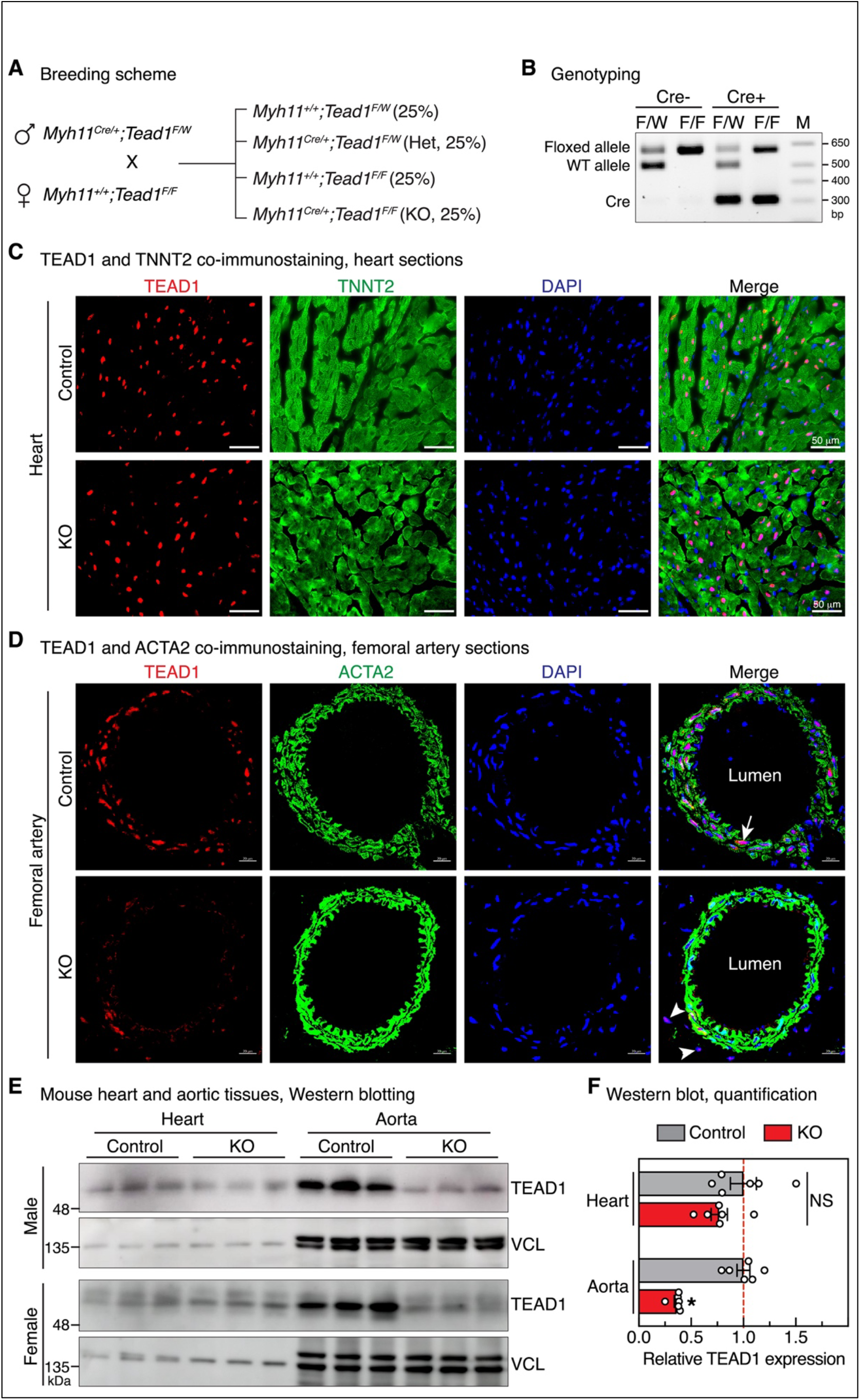
Generation and validation of constitutive SMC-specific *Tead1* KO mice. **A**. Breeding scheme for generating constitutive SMC-specific *Tead1* KO mice. **B**. Representative gel picture of genotyping results using tail biopsy collected from the offspring of the breeding pairs described in “**A**”. M: DNA marker. **C-D**. Heart or femoral artery sections were prepared from control or *Tead1* KO mice. Co-IF assays were then performed to examine the expression of TEAD1 (red) or cardiomyocyte marker TNNT2 (green) (**C**) or smooth muscle marker ACTA2 (green) (**D**) in control or KO mice. DAPI was used to stain nuclei (blue). Arrow and arrowheads denote SMCs- and fibroblast cells-expressed TEAD1, respectively. **E**. Protein lysate was prepared from both male and female control or *Tead1* KO hearts or aorta. Western blotting was then performed to assess TEAD1 expression as indicated. VCL (vinculin) serves as the loading control. **F**. Band densities shown in panel **E** were quantified and plotted. After normalization to the loading control VCL, the relative expression of TEAD1 in control mice was set to 1. N=6. NS: not statistically significant; *p<0.05.

## DISSCUSSION

In this study, we present a new *Myh11*-Cre driver that constitutively expresses Cre recombinase with high specificity and efficiency from the endogenous *Myh11* gene locus. This Cre driver can be used to investigate SMC fate changes during embryogenesis and disease progression by crossing it with reporter mice, such as the mTmG or tdTomato mice as presented here. Furthermore, recent genomic studies have identified many previously unexplored SMC-enriched or -specific genes harboring risk mutations for vascular diseases ^23,24 39^. Our novel *Myh11*-Cre mouse model provides a powerful tool to facilitate the inactivation, activation, or mutation of these genes selectively in SMCs of mice, enabling the testing of causal roles of these risk genes in vascular disease contexts.

The assembly of the vessel wall is a crucial process in the development of embryonic cardiovascular system. Following the formation of the primary vascular endothelial plexus, primordial vascular SMCs are recruited to the endothelium to construct a multilayered vessel wall in a highly organized manner ^40^. Defects in molecular pathways, such as TGF-β signaling, which govern the development and maturation of VSMCs during embryogenesis, can lead to congenital cardiovascular malformations or alter VSMC properties, thereby predisposing the vessel wall to vascular diseases ^41^. A Cre driver that constitutively expresses Cre recombinase driven by *Talgn* promoter has been widely used to investigate SMC biology during embryonic development ^33^. However, a notorious caveat of this Cre line is the transient expression of *Talgn* in cardiomyocytes between E8.5 and E12.5 ^42^. Gene deletion using *Tagln*-Cre often leads to embryonic lethality in mice, displaying both cardiac and VSMC abnormalities as shown with *Tead1* ^1,6,43^. This limitation has considerably hampered efforts to uncover the true role of the gene of interest in VSMC development since cardiac dysfunction may impact VSMC development as a secondary effect, and the appearance of VSMC defects may require an extended time before embryonic lethality caused by lethal cardiac phenotypes. In contrast, *Myh11* expression is well known to be exclusively restricted to SMCs beginning at E10.5 ^29,36^. Therefore, our novel *Myh11*^*Cre*^ KI mouse can be used to circumvent the cardiac phenotype observed after gene deletion with the existing *Talgn*-Cre drivers for developmental studies.

In addition to its transient activation in cardiomyocytes, *Tagln* is expressed in several other cell populations, including myofibroblasts ^44^, myeloid cells ^45^, megakaryocytes and platelets ^46^, vascular progenitor endothelial colony-forming cells ^47^ and perivascular adipose precursor cells ^48^. Given the pivotal roles of these cell types in cardiovascular pathophysiology ^26^, the non-SMC expression of Cre in *Talgn*-Cre drivers introduces confounding effects, largely limiting their utility in studying the sole role of SMCs during disease progression. Therefore, it is conceivable that previous studies showing that deletion of genes using *Talgn*-Cre driver increases susceptibility to vascular diseases such as blood pressure disorder, aneurysm and neointima formation may due to the combinational consequence of gene deletion in multiple cell types ^9-13^. We anticipate that the novel *Myh11*^*Cre*^ KI mouse model reported here will facilitate the dissection of intrinsic effects of losing genes of interest solely in SMCs, eliminating confounding influences from other cell types.

Additionally, most existing SMC Cre drivers are generated through transgenic approaches, raising another concern related to the random (and unmapped) sites of integration plus the variable number of transgene copies ^34^. The traditional transgenic strategy potentially results in unpredictive expression in other cell types ^30,49-51^, aberrant expression of endogenous genes at the integration site by disrupting of critical coding or noncoding sequences such as noncoding RNAs, enhancers and silencers ^32^, introducing extra genes from the transgene if using a large DNA template such as bacterial artificial chromosome ^52^, and ectopic expression of Cre recombinase ^32^. For this novel Cre driver, we used homologous recombination for gene targeting to ensure a single copy of Cre recombinase is expressed from the endogenous *Myh11* gene locus, thus minimizing the potential issues observed with unmapped transgenic Cre drivers.

One caveat of our Cre driver is that the expression of endogenous *Myh11* is disrupted in the KI allele. However, this concern is mitigated as the other WT allele appears to largely compensate the dosage of *Myh11* expression from the loss of the KI allele. Also, maintaining male Cre mice in a heterozygous manner is recommended as a common practice ^26^. It should be noted that mice carrying one Cre allele, without or with one allele of a floxed gene, should be used as controls when utilizing this Cre driver to account for potential haploinsufficiency effects of endogenous *Myh11* gene, and expression of Cre recombinase. Furthermore, *Myh11* is pan-SMC maker expressing in both vascular and visceral SMCs. Gene deletion using Cre lines driven by *Myh11* promoter may lead to visceral myopathies that prevents studying vascular phenotypes. A recently developed *Itga8*-Cre driver exhibits preferential Cre activity in VSMCs with minimal activity in visceral SMCs, offering a valuable tool for studying VSMC function while circumventing visceral SMC phenotypes ^53^. Leveraging single-cell RNA-seq (scRNA-seq) datasets generated in different SMC-enriched tissues may lead to the identification of additional candidates for the development of tissue-restricted SMC Cre drivers.

In summary, we have generated and validated a novel constitutive *Myh11*-Cre mouse model. Alongside recently developed Cre drivers using a similar KI strategy at the *Myh11* gene locus ^31,54^, these new genetic tools are expected to significantly enhance our ability to trace SMC lineage and manipulate gene function specifically in SMCs of both sexes in mice.

## METHODS

### Generation of *Myh11*^*Cre*^ KI mice and *Tead1* SMC-specific KO mice

All mouse studies were approved by Institutional Animal Care and Use Committees at Augusta University (IACUC number: 2012-0502). The *Myh11*^*Cre*^ mice were generated using homologous recombination as we recently reported ^35,36^. An NLS-Cre-ER^T2^-polyA cassette (**Figure 1A**) was inserted immediately after the ATG start codon in exon 2 of the *Myh11* gene. The targeted insertion was confirmed by PCR analysis with primers listed in **Supplemental Figure 1B** and Sanger sequencing. Animals were maintained in C57BL/6J background. The floxed *Tead1* mice were generated as we previously reported ^6,55^. Generation of the *Tead1* constitutive SMC-specific KO mice by using *Myh11*^*Cre*^ mice was depicted in **Figure 4A**. The *Myh11*^*Cre*^ KI mouse line will be made available from the corresponding authors upon request and will be deposited to the Jackson Laboratory.

### DNA extraction, genotyping by PCR (polymerase chain reaction)

DNA was extracted from tail biopsy essentially following the manufacturer’s protocol (Viagen Biotech, cat. #: 101-T). Genotyping by PCR (Thermo Fisher Scientific, cat. #: K1082) was performed using the primers listed below as previously described ^6,55,56^. PCR products were resolved in an agarose gel and visualized under UV after staining with ethidium bromide. We performed multiplex PCR with primers P1 (5’-ATGTCACTCCTTGCCTGAAGGAT-3’), P2 (5’-TCCCTGAACATGTCCATCAGGTTC-3’), and P3 (5’-CACAAAGAGGAACTTCTCATCATCGC-3’) to determine WT allele (P1/P3,

250 bp) and Cre KI allele (P1/P2, 404 bp) for *Myh11* gene locus. For genotyping *Myh11*^*Cre*^-mediated *Tead1* deletion, we used *Tead1* forward primer (5’-TGCCATCATGCCAAGCTATACTGG-3’), reverse primer (CCTGCTTTATAGTCACAGCAGAGGC) and Cre primers (forward, 5’-GCGGTCTGGCAGTAAAAACTATC-3’; reverse, 5’-CTGATCCTGGCAATTTCGGCTATACG-3’) to determine *Tead1* floxed allele (604bp), WT allele (498bp) and Cre (301bp), respectively, as we previously reported ^6^.

### Protein extraction and Western blotting

Aortic and GI tissues were isolated from 8-week-old male or female *Myh11*^*Cre/+*^ mice and their WT littermates (*Myh11*^*+/+*^) obtained by breeding the heterozygous male mice with WT female mice. To determine the specificity and efficiency of *Tead1* deletion in aorta vs heart, heart and aorta were harvested from 10-week-old male and female *Tead1* KO mice. Protein lysate from Cre negative mice (*Myh11*^*+/+*^*;Tead1*^*F/W*^ or *Myh11*^*+/+*^*;Tead1*^*F/F*^) served as controls. Protein lysates were extracted by RIPA buffer (Thermo Fisher Scientific) plus 1% protease/phosphatase inhibitor cocktail (Thermo Fisher Scientific). After sonication and centrifugation of the lysed tissue, proteins in the supernatant were quantified by BCA assay (Thermo Fisher Scientific) and resolved on a 7.5% or 9% SDS-PAGE gel with 10 μg per lane where appropriate. Primary antibodies used in this study were MYH11 (Biomedical Technologies, BT-562, rabbit, 1:1000), Cre (Millipore Sigma, 69050, rabbit, 1:1000), tubulin (TUBA; Cell signaling, 2144, rabbit, 1:1000), TEAD1 (Abcam, ab133533, rabbit, 1:1000), and vinculin (VCL; Sigma, V4505, mouse,1:5000). Secondary antibodies conjugated with horseradish peroxidase were then used to visualize the target proteins in blot. Image was acquired by ImageQuant 800 biomolecular imager (GE) or ChemiDoc™ Imaging System (Bio-Rad) and band densities were quantified using ImageJ.

### Sections and GFP visualization

To verify Cre recombinase expression and activity, *Myh11*-Cre male mice were first crossed with the mTmG female reporter mice (Jackson Laboratory; B6.129 (Cg)-Gt(ROSA)26Sor^tm4(ACTB-tdTomato,-EGFP)Luo^/J; Stock No: 007676). Aortic tissues and other tissues from 8-week-old male *Myh11*^*Cre/+*^*;mTmG*^*F/W*^ mice and their WT littermates (*Myh11*^*+/+*^*;mTmG*^*F/W*^) were harvested, photographed under bright field or epifluorescence GFP channel, and subsequently fixed with 4% paraformaldehyde (PFA) on ice for 30 minutes. After washing with ice-cold phosphate-buffered saline (PBS) twice for 10 minutes each time, tissues were then soaked in 30% sucrose/PBS on a shaker in 4°C overnight until the tissues settled down. The processed tissues were subsequently embedded in optimal cutting temperature (OCT) medium (Tissue-Tek) and cryosections were prepared at 8-μm thickness. For directly visualizing GFP, OCT sections were air dried for 10-15 minutes then washed with PBS (3 times x 5 minutes/wash) to remove OCT. The slides were then briefly air dried and covered with Prolong Gold DAPI mounting medium (Invitrogen, P36962) to counter-stain nuclei. GFP and DAPI signals were then acquired by a confocal microscopy (LSM 780 upright, Zeiss) using 555-nm (tdTomato), 488-nm (EGFP), 405-nm (DAPI) excitation channels, respectively.

### RFP visualization and co-immunofluorescence (co-IF) staining

To verify whether Cre recombinase expression and activity overlap the expression of endogenous *Myh11* gene, *Myh11*-Cre male mice were crossed with the tdTomato female single fluorescence reporter mice (Jackson Laboratory; B6.Cg-Gt(ROSA)26Sor^tm9(CAG-tdTomato)Hze^/J; Stock No: 007909) and co-IF was performed on their offspring *Myh11*^*Cre/+*^*;tdTomato*^*F/W*^ mice. Embryonic day (E) 0.5 was defined as noon of the day when the vaginal plug was detected. Embryonic or adult mouse tissues were collected in PBS, photographed under bright field or epifluorescence RFP channel, and subsequently fixed in 4% PFA on ice for 20 minutes to 1 hour according to the developmental stage. After washed in PBS three times the tissues were dehydrated with 30% sucrose overnight at 4^°^ C before being embedded in OCT compound on dry ice. Embedded samples were then sectioned at 10 microns with a cryotome (Leica, CM3050S). Sections were blocked in blocking solution containing 10% donkey serum (Sigma, D9663) and 0.1% Triton in PBS (PBST) for 1 hour at room temperature, followed by incubation with anti-MYH11 primary antibody (Abcam, ab53219, rabbit, 1:50) and anti-PECAM1 primary antibody (R&D Systems, AF3628, goat, 1:50) overnight at 4^°^ C. After three washes with 0.1% PBST, sections were incubated with Donkey anti-Rabbit secondary antibody conjugated with Alexa Fluor™ Plus 488 (Invitrogen, A32790, 1:200; For detecting MYH11) and Donkey anti-Goat secondary antibody conjugated with Alexa Fluor™ Plus 647 (Invitrogen, A32849, 1:200; For detecting PECAM1) for 2 hours at room temperature. Nuclei were stained with Hoechst 33342 (Invitrogen, H3570, 1:10000) for 5 minutes. Images were obtained using a Leica fluorescence microscope with 405-nm (Hoechst 33342), 488-nm (MYH11), 555-nm (native tdTomato signal from the reporter) and 647-nm (PECAM1) excitation channels, respectively.

### Determining TEAD1 expression by tyramide signal amplification (TSA) staining

For IF staining of TEAD1 in the heart and femoral artery, TSA kit (TSA plus DIG, Perkin Elmer, cat. #: NEL748001KT) was used to amplify the TEAD1 signal as we previously reported ^56^. Briefly, after antigen retrieval as done by using microwave to heat at 98°C for 10 minutes in citric acid buffer (10mM, pH6.0), sections were treated with H_2_O_2_ (30 uL 30% H_2_O_2_ diluted into 1 mL PBS) for 15 minutes and then incubated with blocking buffer (2.5% horse serum + 0.1% Tween-20) for 30 minutes. Following immunostaining with anti-TNNT2 (Cardiac troponin T2; DSHB, RV-C2; Mouse; 1:50) or ACTA2 (SM α-actin; Sigma, A2547, mouse, 1:200) antibodies, anti-TEAD1 antibody (Abcam, ab133533, rabbit, 1:250) was incubated with the sections overnight at 4°C. After incubation with HRP labeled anti-rabbit secondary antibody (Vector Labs, cat. #: MP-7401) for 30 minutes at room temperature, TSA plus DIG Amplification working solution (1:50 dilution) was applied for 5 minutes at room temperature. Sections were then further incubated with cy5 fluorescence-labeled anti-DIG antibody (TSA Plus Cyanine 5, NEL745E001KT, 1:250 dilution) and Alexa Fluor™ 555 (Invitrogen, Goat anti mouse, A-21422, 1:250) for 30 minutes at room temperature. Finally, sections were mounted with Prolong Gold mounting medium with DAPI (Invitrogen, P36962) and images were acquired by a fluorescence microscopy (REVOLVE, ECHO) or a confocal microscopy (LSM 780 upright, Zeiss).

### Statistical Analysis

GraphPad Prism (version 9.2) was used for the statistical analysis. All data are expressed as mean±SEM of at least 3 independent experiments. An unpaired 2-tailed *t* test was used for data involving 2 groups only. Values of *P*<0.05 were considered statistically significant.

## SUPPLEMENTAL FIGURES

**Supplemental Figure 1.**
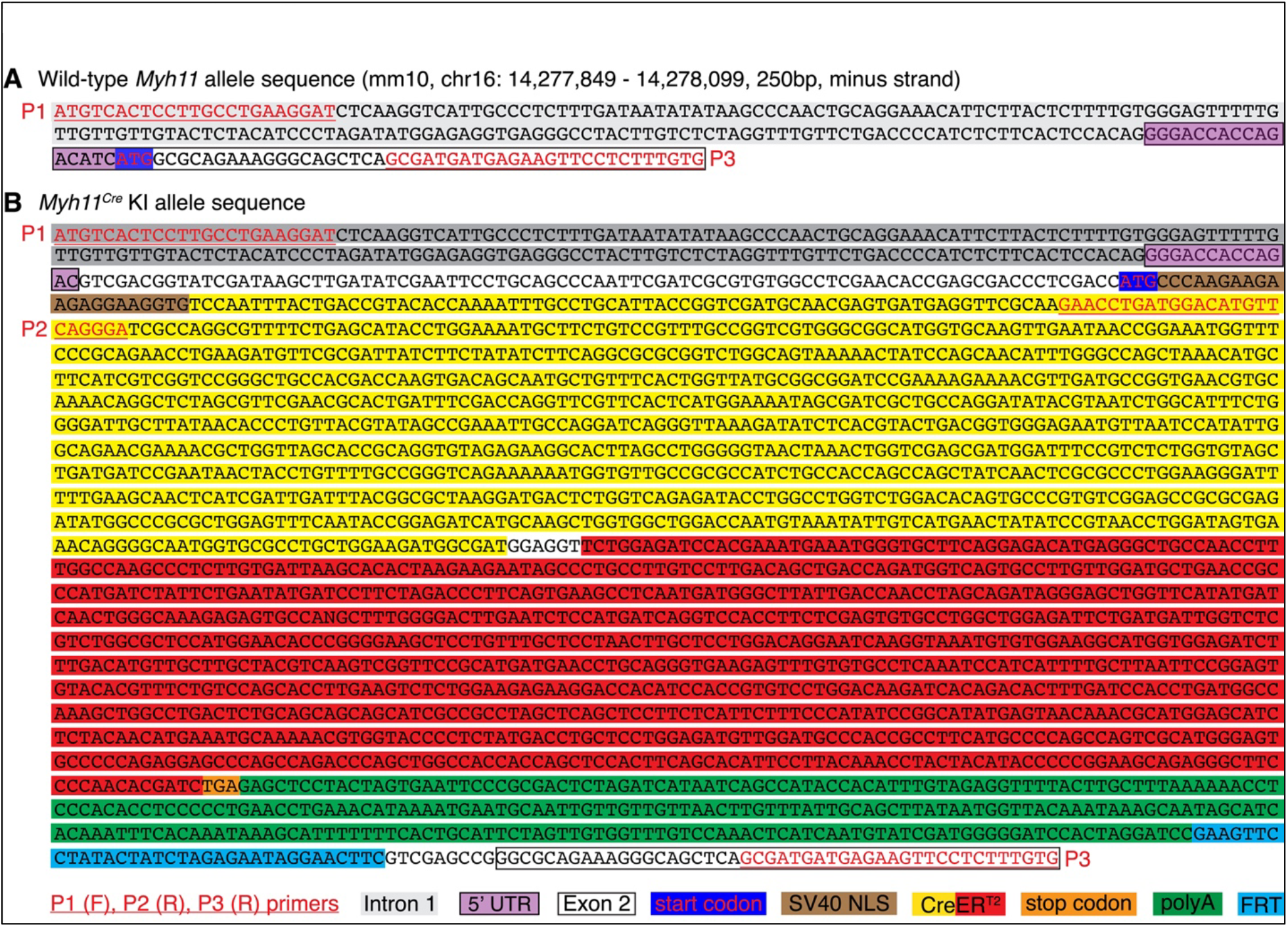
Validation of the *Myh11*^*Cre/+*^ KI allele by DNA sequencing. Nucleotide sequence of the P1/P3 PCR amplicon using primers to the *Myh11* gene locus of the WT allele (**A**) or *Myh11*^*Cre*^ KI allele (**B**), respectively. P: primer; F: forward; R: reverse.

**Supplemental Figure 2.**
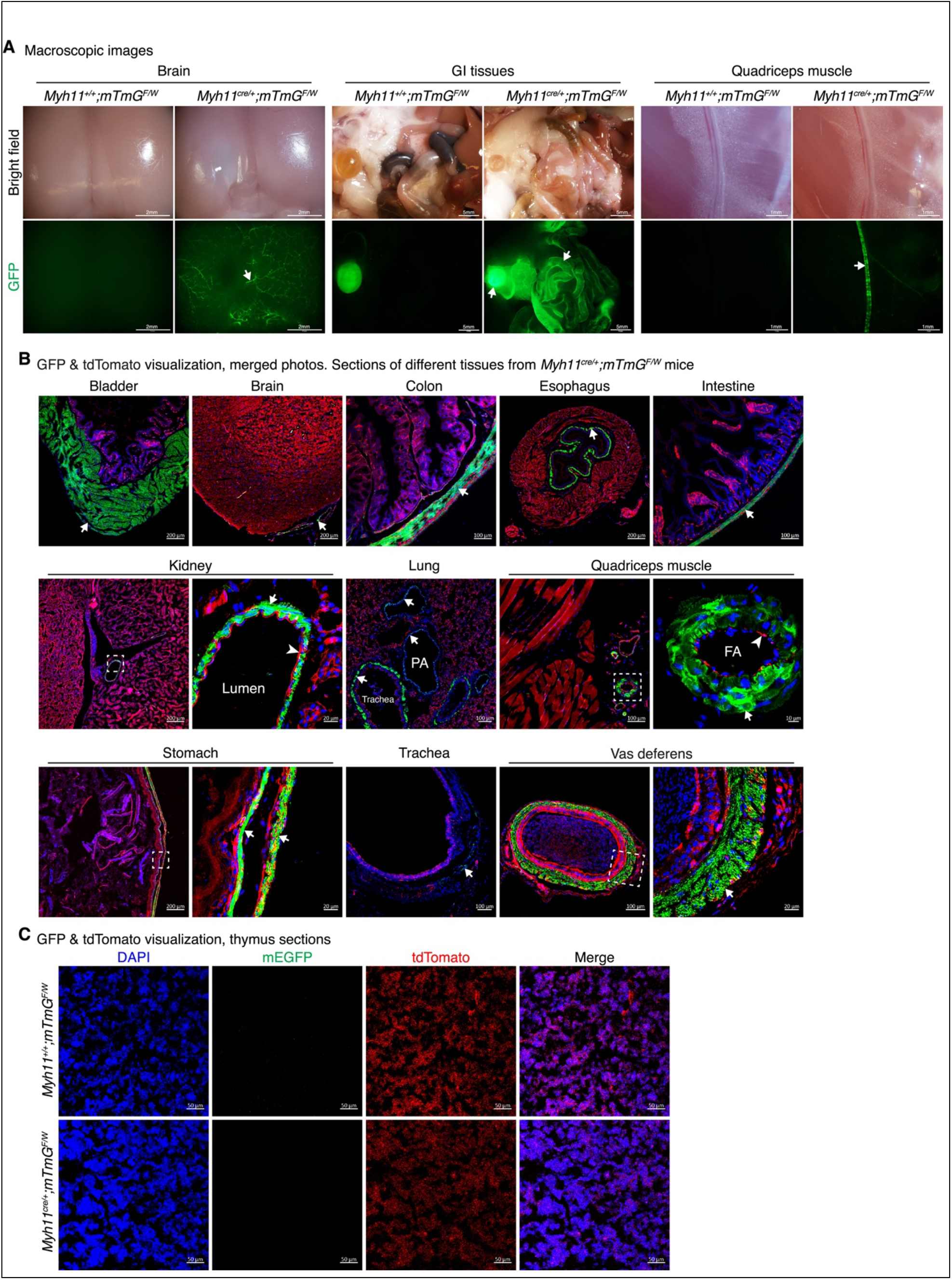
Cre activity of the *Myh11*-Cre driver in other tissues. **A.** Macroscopic images were acquired with brain, gastrointestinal (GI) tissues and quadricep muscle of control and mTmG SMC lineage tracing mice under bright field (top panels) or GFP channel (bottom panels). Arrows point to GFP positive area including microvessels on the surface of the brain (left), bladder and small intestine of the GI tracts (middle) and femoral artery on the surface of quadricep muscle (right panel). **B**. mEGFP (mG, green) and tdTomato (mT, red) visualization in sections of bladder, brain, colon, esophagus, intestine, kidney, lung, quadriceps muscle, stomach, trachea and vas deferens of *Myh11*^*Cre/+*^*;mTmG*^*F/W*^ SMC lineage tracing mice. Cell nuclei were counter stained by DAPI (blue). Arrows point to Cre-dependent GFP signals. Arrowheads indicate tdTomato positive endothelial cells. The boxed area is magnified to the right. FA: femoral artery. **C**. Thymus was collected from adult control (top) and SMC lineage tracing mice (bottom panel) for direct visualization of mEGFP (green) and tdTomato (red) signals. DAPI was used to counter stain cell nuclei (blue).

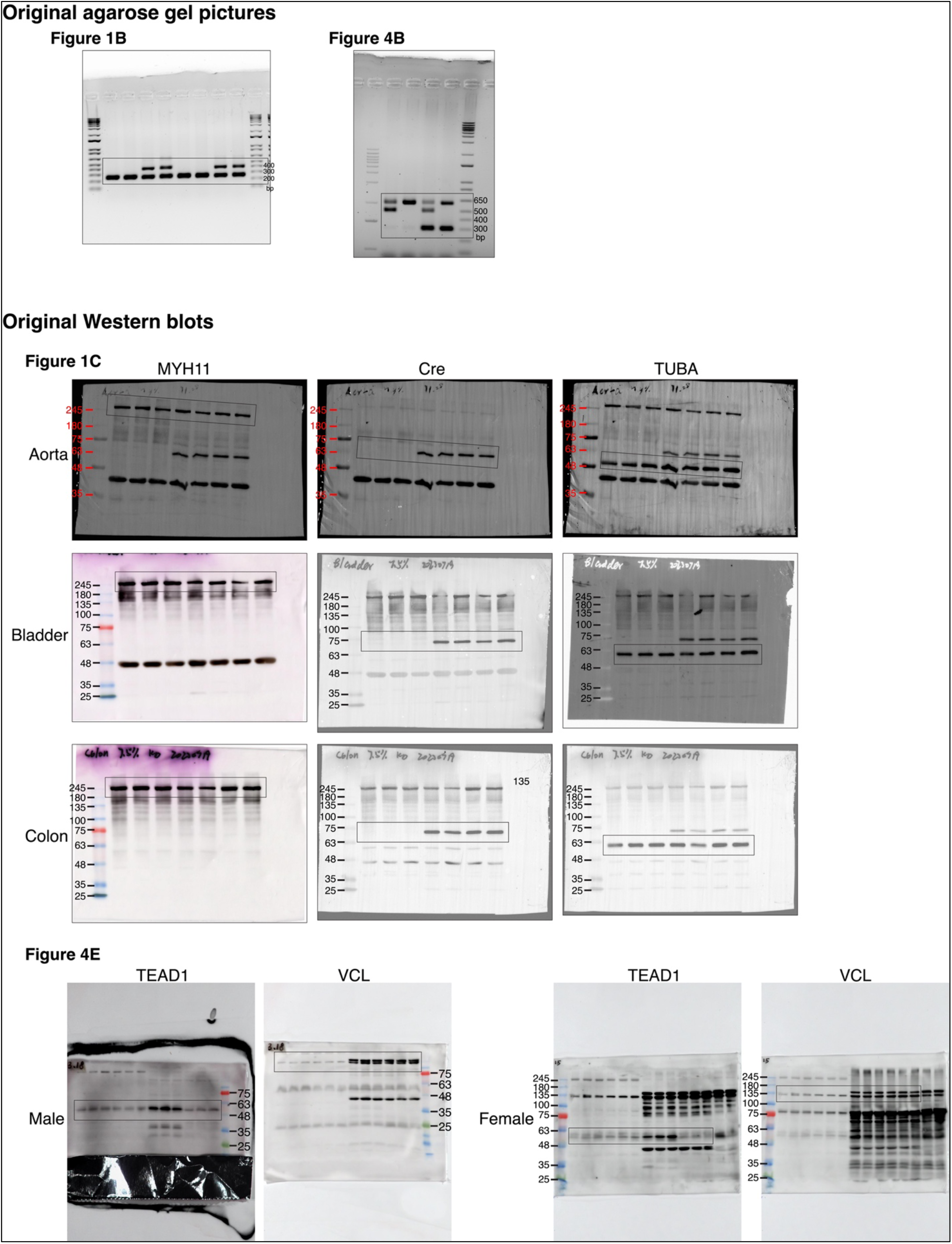

**Original agarose gel pictures and Western blots**

## Notes

**Conflicts of Interest:** The authors disclose no conflicts.

**Funding:** The work in Jiliang Zhou’s laboratory is supported by grants from the National Institutes of Health and the National Heart, Lung, and Blood Institute (HL157568 and HL149995). Kunzhe Dong and Xiangqin He are supported by a Career Development Award (938570) from the American Heart Association and a K99 award (K99HL171834) from the NIH, respectively.

### Competing Interest Statement

The authors have declared no competing interest.

